# Exploring the Restoration of Memory Post-Neuronal Regeneration from Induced ALS-FTD in *Dugesia tigrina*

**DOI:** 10.1101/2025.10.30.685683

**Authors:** Arsh Jha, Janani Ramkumar, Aarush Jugdar, Xiuxia Du

## Abstract

Amyotrophic lateral sclerosis–frontotemporal dementia (ALS–FTD) is a neurodegenerative disease characterized by progressive neuronal loss and motor decline. Recovery remains beyond therapeutic reach, and even complete neurogenesis does not ensure preservation of memory capacity (defined as the strength of synaptic connections). Freshwater planarians, capable of whole-body regeneration and associative learning, provide a tractable model to study the relationship between neuronal damage and behavior. Their “single-circuit” centralized nervous systems allow evaluation of oxidative stress, a key driver of ALS–FTD pathology, which was induced *via* hydrogen peroxide.

In this study, 180 planaria underwent three phases. Phase 1 established baseline memory using a 28-day operant conditioning protocol with a light-based training device. Success was determined by the proportion of planaria in concentric circles defined as zones (A-C) with a respective gradient descent of light concentration. Phase 2 determined hydrogen peroxide (H_2_O_2_) concentrations sufficient to induce oxidative stress that triggered regenerative signaling and neuronal damage without lethality. Phase 3 applied the optimal concentration, allowed regeneration, and retested memory.

The optimal concentration of hydrogen peroxide was 5 ppm, triggering blastema formation and neuronal damage while preserving viability. Zone occupancy analyses utilizing prior found concentration revealed measurable post-exposure shifts: Zone A increased initially but trended lower post-exposure, Zone B increased, and Zone C decreased significantly. These results indicate that while memory capability persists after neurogenesis, the quality of memory decreases. In accordance with this study, current treatments for ALS–FTD that focus solely on neurogenesis will not fully restore cognitive function.

## 1. Introduction

ALS-FTD (amyotrophic lateral sclerosis–frontotemporal dementia) involves progressive neuronal loss that impairs both motor and cognitive function. Current treatments cannot restore lost neurons, and even if complete neurogenesis were achieved, it remains unclear whether structural repair alone would be sufficient to preserve previously acquired behaviors and memories (Chung et al., 2022). This raises an important question: after full neuronal regeneration in a small, defined closed system cluster of neurons (circuit), do the regenerated neurons possess the same quantitative capacity for memory as before injury? Addressing this question requires a model system capable of both neural regeneration and measurable behavior.

Freshwater planarians (Dugesia tigrina) are well-established model organisms in regenerative biology and neurobiology due to their remarkable capacity for whole-body regeneration, supported by pluripotent adult stem cells known as neoblasts (Rossi et al., 2022). Their relatively simple but centralized nervous system, bilateral symmetry, and sensitivity to environmental perturbations make them uniquely suited for linking regeneration with behavior (Reddien, 2004). Planarians are capable of both classical and operant conditioning, forming associations between stimuli such as light, vibration, and chemical cues with food reward (Deochand et al., 2018). They also display innate behaviors such as negative phototaxis, the tendency to avoid illuminated areas, which can be overridden through operant conditioning when planarians are reinforced for entering illuminated zones despite their innate avoidance to light (Paskin et al., 2014). Notably, memory traces in planarians can persist even after surgical removal and regeneration of the brain, underscoring the robustness of memory consolidation in this model (Blackiston et al., 2015).

Together, these features make planarians uniquely valuable for examining how neural injury and regeneration influence learning and memory.

Within ALS–FTD, neuronal degeneration is strongly linked to oxidative stress caused by reactive oxygen species (ROS). Hydrogen peroxide (H_2_O_2_) is one such ROS that, while acting as a signaling molecule at low levels, becomes toxic at higher concentrations, damaging DNA, proteins, and lipids, and thereby reducing cell viability (Chung et al., 2022). In planarians, H_2_O_2_ exposure disrupts regeneration by inhibiting neoblast proliferation, inducing apoptosis, and interfering with blastema formation (Pirotte et al., 2015; Wu & Li, 2018). To simulate the effects of oxidative stress, we exposed planarians to carefully titrated concentrations of H_2_O_2_. First, their conditioned memory performance was measured under normal conditions as a baseline control. They were then exposed to an experimentally determined concentration of H_2_O_2_, as no prior studies provided a standardized dosage, and their post-exposure memory performance was compared with baseline. We hypothesized that acute H_2_O_2_ exposure would reduce conditioned performance and that both groups would show behavioral decline relative to baseline.

In this study, we worked with a fresh cohort of 180 D. tigrina planarians, randomly assigned into two experimental groups of 30 each: one conditioned before hydrogen peroxide exposure (pre-H_2_O_2_) and one conditioned after exposure (post-H_2_O_2_). Both groups were trained twice per week to enter an illuminated zone to obtain food, thereby operantly conditioning them to overcome negative phototaxis, and performance was tested in probe trials conducted without reinforcement. In Phase 2, we established that the selected concentration of H_2_O_2_ produced consistent morphological and behavioral stress while maintaining survival. This design allowed us to directly evaluate how oxidative stress influences both the establishment and the expression of conditioned behavior by comparing planarians trained before oxidative stress with those trained afterward.

Unlike prior studies that examine whole-brain regeneration or generalized behavioral recovery, our approach targets a defined subset of neurons within a single circuit, allowing us to isolate how localized damage and regeneration affect memory retention. Planarians are uniquely suited for this because their nervous systems are non-redundant and anatomically simple, enabling precise behavioral readouts from small-scale neural changes. Importantly, planarian regeneration does not result in circuit duplication or reassignment, the original circuit is reconstituted within the same anatomical context, allowing us to ask whether memory traces encoded in a few neurons can survive oxidative damage and be functionally restored. This contrasts with neurodegenerative models like ALS–FTD, where widespread neuronal loss obscures the contribution of individual circuits to cognitive decline. By focusing on a minimal regenerative unit, we aim to clarify whether structural repair within a circuit is sufficient to restore learned behavior.

## 2. Materials and Methods

### 2.1 Animals and Housing

Brown planaria (Dugesia tigrina) were obtained from Carolina Biological Supply (Burlington, NC, USA) and maintained in Poland Spring water at 20–22 °C on a 12 h:12 h light–dark cycle. Water was changed weekly, and animals were fed a pea-sized portion of hard-boiled egg yolk (∼0.25 mL per dish) twice weekly. After feeding, planaria were transferred to clean dishes with fresh spring water to minimize fouling and were starved for at least 48 h before behavioral testing to reduce variability in activity levels.

Only intact adult worms (8–12 mm length) exhibiting smooth gliding behavior without frequent reversals or erratic contractions were used.

### 2.2 Quality Control

A comprehensive quality control protocol was implemented to ensure accuracy, reproducibility, and reliability of all experimental procedures. Before each session, students serving as biology and chemistry teaching assistants cleaned all work surfaces with laboratory-grade disinfectant and confirmed that instruments and reagents were in proper working condition.

All glassware, including beakers, graduated cylinders, and Petri dishes, was washed with laboratory detergent, rinsed thoroughly with deionized water, and dried with KimWipes prior to use. Pipettes (Eppendorf Research Plus) were verified for volume accuracy using distilled water and recalibrated when necessary. Reagents were inspected for clarity and expiration, and the 30% hydrogen peroxide stock solution (Sigma-Aldrich) was stored in opaque containers to prevent light degradation. Working dilutions were prepared fresh before each trial to minimize decomposition and ensure consistent concentration.

The stereomicroscope used for behavioral and morphological observations was cleaned daily with lens paper and ethanol solution to maintain optical clarity. The light source and digital timer were tested before each use to verify illumination stability and timing accuracy. Grid paper beneath the assay dishes was replaced regularly to maintain consistency in spatial measurements across replicates.

Planarian maintenance was also subject to daily quality checks. All animals were kept in Poland Spring water verified to be free of chlorine, heavy metals, and visible particulates. Water quality was assessed weekly for pH and temperature, which were maintained between 6.8 and 7.4 and at 20 to 22 °C, respectively. These parameters were selected to support optimal survival, regeneration, and behavioral stability. Housing dishes were inspected daily for fouling or microbial contamination and replaced immediately when necessary. Only intact adult *Dugesia tigrina* (8–12 mm) displaying smooth gliding movement ad no signs of injury were selected for behavioral trials.

Feeding schedules and starvation periods followed the procedure described in Section 2.1 to reduce metabolic variability. After feeding, planaria were transferred to clean dishes containing fresh spring water to minimize waste accumulation. During the study, environmental factors such as light exposure, vibration, and humidity were monitored to maintain consistent experimental conditions.

All cleaning, calibration, and inspection procedures were documented in the laboratory notebook to confirm compliance and to ensure reproducibility across all experimental phases.

### 2.3 Experimental Apparatus

Behavioral assays were conducted in sterile 90-mm Petri dishes (Corning Inc., Corning, NY, USA) containing 2–3 mm of spring water. A localized light source was positioned above each dish to create three concentric zones of equal radial width (∼10 mm each): the illuminated central zone (Zone A), the surrounding annulus (Zone B), and the outer ring adjacent to the dish wall (Zone C). Grid paper placed beneath the dishes ensured consistent placement and facilitated quantitative scoring. All behavioral trials were conducted in a vibration- and noise-free environment.

### 2.4 Experimental Design

The study was divided into three sequential phases. Phase 1 established a baseline memory curve by training planaria in light–place conditioning and measuring conditioned preference. Phase 2 determined the hydrogen peroxide (H_2_O_2_) concentration that reliably induced acute oxidative stress while preserving survival. Phase 3 applied this concentration to a separate group of planaria and repeated the Phase 1 conditioning protocol, enabling direct comparison of pre- and post-injury learning and memory performance.

At the start of the experiment, 180 planaria were pooled and randomly assigned into six replicate dishes of 30 worms each. Three dishes were allocated to Phase 1 (pre-H_2_O_2_ memory testing), while three dishes were reserved for Phase 3 (post-H_2_O_2_ exposure and recovery). Randomization ensured that observed differences between groups reflected treatment effects rather than pre-existing variation in worm behavior or morphology.

### 2.5 Phase 1: Training Protocol and Probe Sessions

In Phase 1, planaria underwent associative conditioning to confirm their ability to form a learned preference under the assay conditions. Training was performed twice weekly for four weeks (8 sessions total). Each 30-minute session paired localized light in Zone A with food delivery, creating a positive reinforcement association between light exposure and feeding.

Non-reinforced probe sessions were conducted immediately after training sessions to evaluate memory. Thirteen probes were carried out on Days 0, 2, 5, 7, 9, 12, 14, 16, 18, 20, 21, 26, and 28, with standardized benchmarks at Days 7, 14, 21, and 28 to assess acquisition and retention of conditioned responses. The twice-weekly training schedule over four weeks was selected based on prior operant conditioning studies in planaria, which demonstrate that behavioral acquisition and memory stabilization require extended, spaced exposure to reinforcement stimuli (Krantz, 1970; Abbott & Wong, 2008). Probe sessions were distributed across 28 days with standardized benchmarks to capture both early learning and long-term retention, consistent with protocols used in directional bias and maze learning assays.

Distributed training has been shown to enhance long-term memory formation across taxa, including invertebrates, due to the spacing effect (Cepeda et al., 2006). In planarian models, spaced reinforcement avoids overstimulation and supports neurophysiological consolidation (Deochand et al., 2018), which is particularly relevant given their regenerative capacity and neuroplasticity.

### 2.6 Probe Procedure

Each probe session lasted 10 minutes and was performed without food reinforcement to ensure that behavior reflected conditioned memory rather than immediate reinforcement. At the start of every probe, worms were gently transferred to Zone C to standardize initial position. The number of worms in Zones A, B, and C was recorded once per minute, yielding a 10-point time series for each dish. Counts were normalized to proportions (number in zone ÷ total worms in dish), producing standardized memory scores across replicates.

### 2.7 Phase 2: Preparation of H_2_O_2_ Working Solutions and Concentration Screen

To determine a reproducible sublethal dose, hydrogen peroxide working solutions were prepared by serial dilution of a 30% stock solution (Sigma-Aldrich, St. Louis, MO, USA) in spring water. A 1000 ppm intermediate solution was generated and diluted to produce 3 ppm and 10 ppm working concentrations. A 2% v/v solution was prepared directly from the stock for comparison. All dilutions were freshly prepared prior to use to minimize decomposition.

Planaria were exposed to each concentration, and morphological as well as behavioral responses were monitored at defined time points (immediate, 1 min, and post-recovery). Morphological changes were quantified by measuring body length and width against 1 cm grid paper. Behavioral responses included presence or absence of erratic movement, duration of erratic movement, directed movement toward or away from light, and responsiveness to mechanical stimulation using a stream of water. Erratic behavior was defined as rapid, uncoordinated locomotion characterized by twisting, spinning, frequent reversals, and vigorous contractions inconsistent with gliding.

Responses were timed from onset until worms either resumed gliding, displayed pulsing recovery movements, or showed signs of imminent mortality such as extreme contraction or visible tissue damage. Blastema formation at 24 h was also recorded as a marker of regenerative viability. From this screen, 5 ppm H_2_O_2_ was identified as the lowest reproducible dose that produced robust stress responses while preserving survival and was selected for Phase 3.

### 2.8 Phase 3: Acute H_2_O_2_ Exposure and Behavioral Assays

At the conclusion of Phase 1 (Day 28), the dishes reserved for Phase 3 were exposed to acute oxidative stress. A 10 µL droplet of freshly prepared 5 ppm H_2_O_2_ was applied directly to the dorsal surface of each worm with a micropipette (Eppendorf Research Plus), immediate responses were noted under a dissecting microscope, and animals were then returned to fresh spring water. Following exposure, worms were maintained under standard housing without behavioral testing for a 14-day recovery period to allow healing and neurogenesis.

After this 14-day recovery, the Phase 1 conditioning protocol was repeated in full. Animals were trained twice per week for four weeks (eight 30-minute sessions) with light in Zone A paired with food delivery, and each training session was followed immediately by a 10-minute non-reinforced probe. Probe scheduling mirrored Phase 1 exactly, with 13 probes on Days 0, 2, 5, 7, 9, 12, 14, 16, 18, 20, 21, 26, and 28. All handling, scoring, and data processing were identical to those described for Phase 1 (Sections 2.4–2.5 and 2.8).

### 2.9 Data Processing

For each probe session, data points were defined by three indices: probe day (y ∈ [0, 28]), zone (n ∈ {A, B, C}), and probe minute (t ∈ [0, 10]). At each minute, the number of planaria in a given zone was normalized to a proportion:

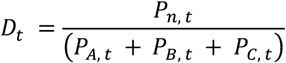

where p denotes planaria, t is time in minutes, n is the zone, and Dt is the data point. The mean over the analysis window was calculated as:

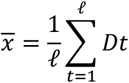

with ℓ the total number of valid samples in the domain Dn(t). For all analyses, minutes 2–10 were included.

2. If variability exceeded a pre-specified threshold (> #), the affected Dt was excluded and the mean recomputed without that point:

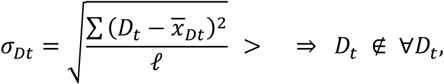

where 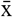 is the mean proportion in zone n and ℓ is the total number of valid samples. Only minutes 2–10 were analyzed to reduce exploratory noise.

If variability exceeded a pre-defined threshold, the affected Dt was excluded and the mean recomputed without that point. Outliers were thus removed, and the final dish-level score was the re-averaged mean of the remaining samples. Zone C was recomputed as C = 1 − A − B to ensure closure. Triplicate dish scores were averaged to produce group means ± SD. Each treatment group (pre-H2O2 and post-H2O2) contained three replicate dishes (30 worms each, 90 total per group). For a given probe day (e.g., Day 7), dish-level means were averaged across replicates to yield the group mean ± standard deviation (SD). These group means formed the data points shown in the zone-occupancy trajectories, with error bars representing variability across replicates.

In this way, every dot on the final graphs corresponds to the mean proportion of worms in a zone on a specific probe day, derived from normalized per-minute counts, averaged first within dishes and then across replicates.

### 2.10 Statistical Analysis & Reproducibility

Statistical analysis was used to determine whether hydrogen peroxide (H2O2) exposure caused measurable changes in planarian learning and memory compared to baseline behavior. Each condition (pre-H2O2 and post-H2O2) included three biological replicates (n = 3 dishes, 30 worms per dish, 90 worms per group).

For every 10-minute probe session, the proportion of worms in Zones A, B, and C was recorded, and the mean values were compared across probe days. All data are reported as mean ± standard deviation (SD).

Welch’s two-sample *t*-test was applied to compare the pre- and post-H2O2 groups for each zone at each probe day. This test was chosen because it allows comparison of group means without assuming equal variances, which was important since the behavioral variability of the worms could differ after oxidative stress. The test helped determine whether the memory performance of planaria changed significantly following H2O2 exposure. Degrees of freedom were calculated using the Welch–Satterthwaite equation, and all tests were two-tailed with the significance level set at α = 0.05.

Effect sizes were estimated using Cohen’s *d* to show the magnitude of any observed differences. Because behavioral probes were conducted repeatedly over time, results were interpreted as part of the overall behavioral pattern rather than as individual time points.

All statistical analyses and figures were generated in Python (v3.11) using the numpy, pandas, scipy, and matplotlib libraries. This approach provided consistent data handling and reproducible calculations across trials.

## Results

The following results describe the effects of acute hydrogen peroxide (H_2_O_2_) exposure on planarian behavior, morphology, regeneration, and conditioned zone occupancy. Data are presented as mean ± SD and include statistical comparisons of pre- and post-exposure conditions. Figures 1 and 2 and Supplementary Tables S1–S3 summarize concentration-dependent behavioral and morphological changes, regenerative outcomes, and shifts in zone occupancy over the 28-day period.

**Figure 1.**
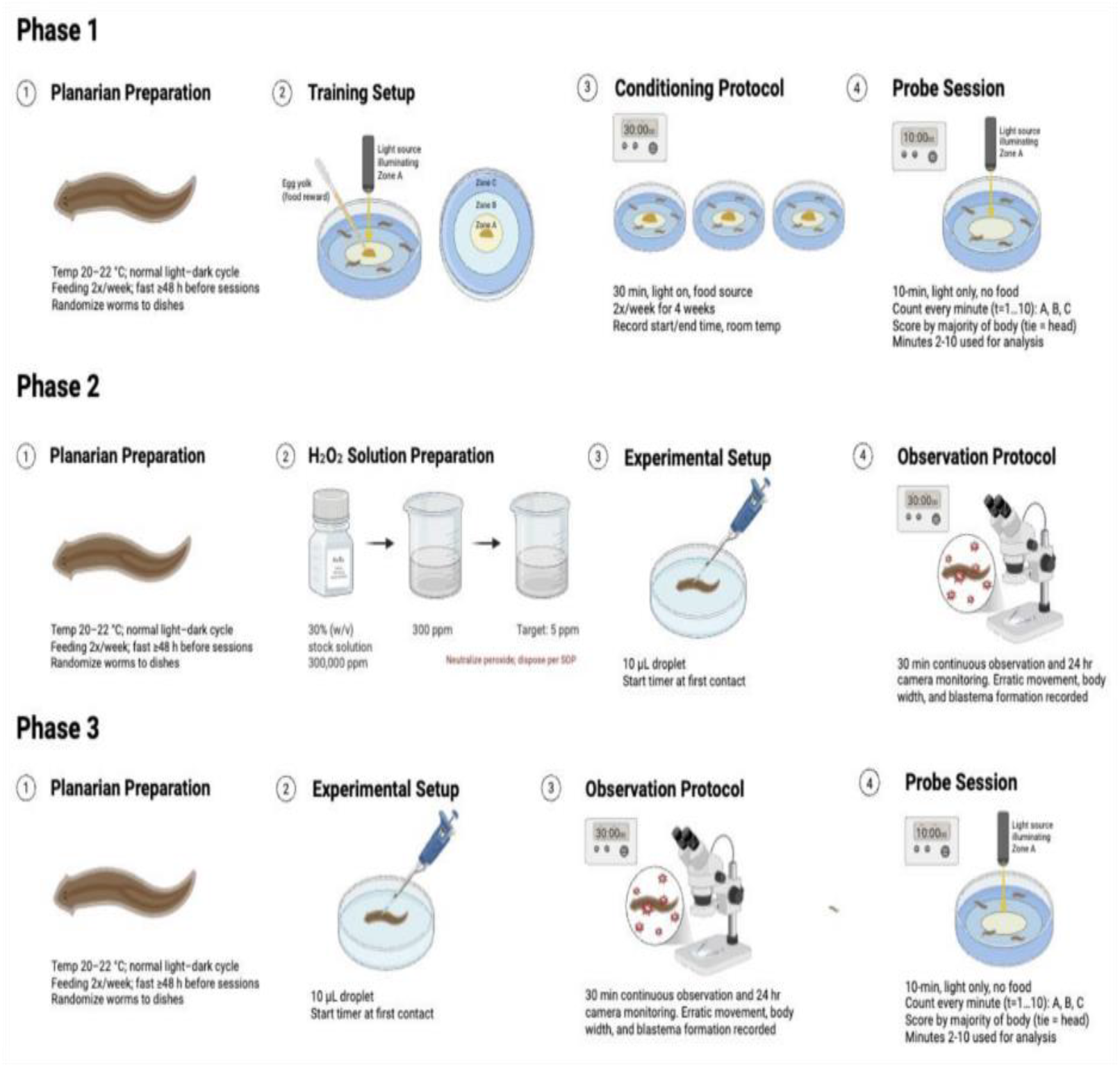
Experimental design for evaluating oxidative stress effects on planarian learning and memory. (a) Phase 1: Planarian preparation, training setup, conditioning protocol, and probe sessions to establish baseline memory through food-reinforced light conditioning. (b) Phase 2: Preparation of H_2_O_2_ working solutions from a 30% stock, experimental setup with dorsal droplet exposure, and observation protocol to monitor erratic movement, body width, and blastema formation. (c) Phase 3: Post-exposure regeneration and retesting using the same conditioning and probe protocols as Phase 1 to compare memory performance before and after oxidative stress.

**Figure 2.**
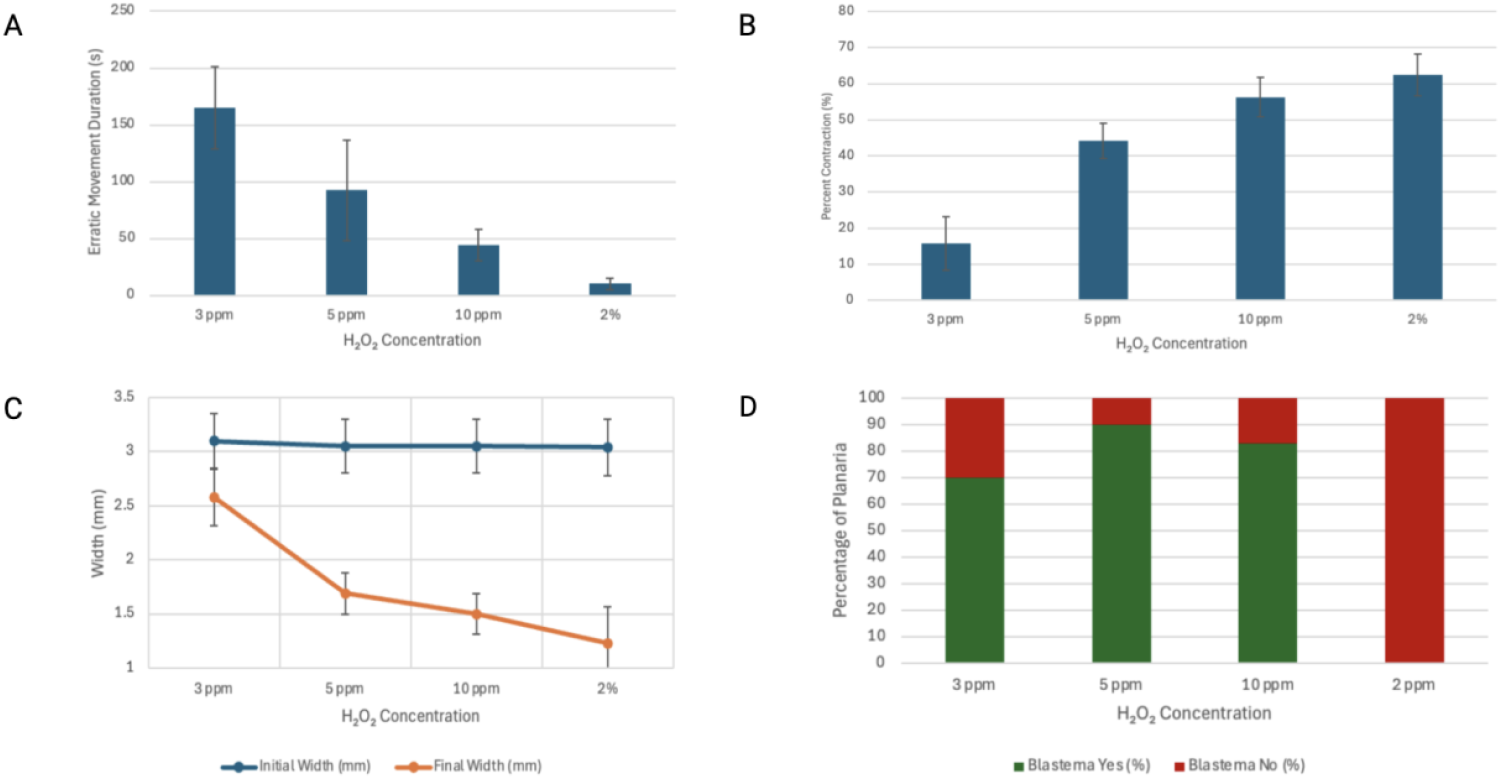
Concentration-dependent effects of hydrogen peroxide (H_2_O_2_) on planarian behavior, morphology, and regeneration. (a) Mean duration of erratic movement (seconds ± SD) for planaria exposed to 3 ppm, 5 ppm, 10 ppm, or 2% H_2_O_2_. Lower concentrations (3 and 5 ppm) induced prolonged movement, whereas higher concentrations (10 ppm and 2%) produced brief spasmodic movements and rapid nonviability. (b) Percent body width contraction, showing increasing contraction with H_2_O_2_ concentration and maximal morphological collapse at 2%. (c) Final body widths (mm ± SD) decreased progressively with increasing concentration. (d) Proportion of planaria exhibiting visible blastema formation at 24 hours post-exposure, with visible stress responses at 5 ppm and absence of blastema at 2%. Data represents n = 30 planaria per concentration.

**Figure 3.**
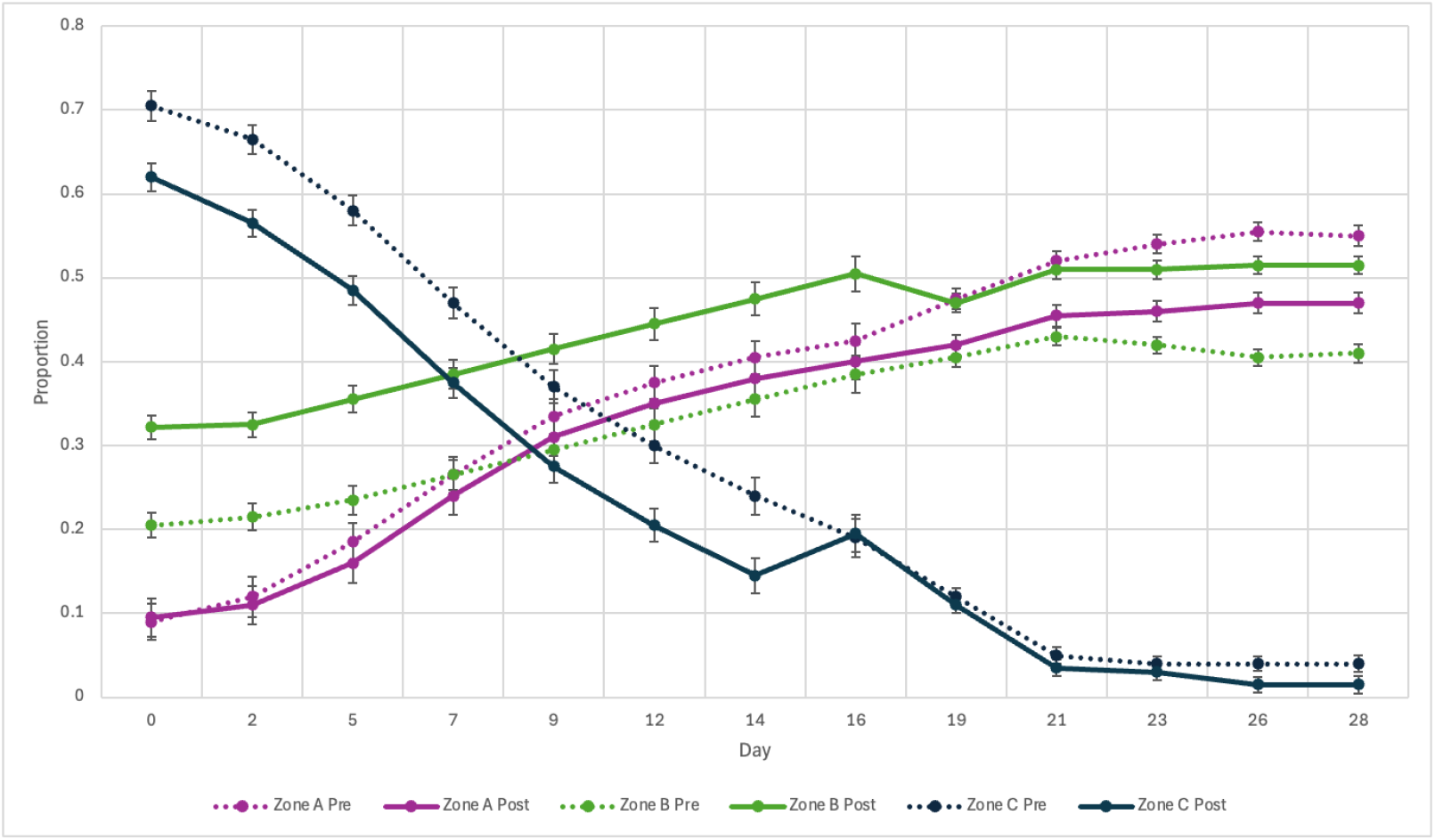
Validation of zone occupancy in planaria under light–place conditioning before and after H_2_O_2_ exposure. Proportion of planaria located in Zones A (center), B (annulus), and C (outer ring) is shown across 28 days (13 probe sessions) for pre-(solid lines) and post-exposure (dashed lines) conditions. Occupancy in Zone A increased gradually over time with post-exposure reductions, Zone B showed consistent post-exposure increases, and Zone C decreased post-exposure, particularly after Day 14. Data represent mean ± SD across n = 5 trials per day.

Exposure of planaria to increasing concentrations of hydrogen peroxide (H_2_O_2_) produced clear, concentration-dependent effects on behavior, morphology, and regenerative outcomes (Figure 1). As shown in panel (a), planaria exposed to 3 ppm exhibited the longest sustained erratic movement, often exceeding 2 minutes, whereas 5 ppm induced a moderate but sharper response with movement lasting ∼60–120 seconds, followed by visible recovery. At 10 ppm, erratic movement was limited to short bursts (∼30–60 seconds) before stillness, and planaria exposed to 2% H_2_O_2_ displayed immediate spasmodic movement for less than 15 seconds before becoming nonviable. Morphological contraction increased with concentration (panel b), with the most significant reduction occurring at 2%, over 50% within seconds, while intermediate contraction was observed at 5 ppm and 10 ppm, and minimal contraction occurred at 3 ppm. Percent contraction ((Initial Width − Final Width)/Initial Width × 100%) reflected a 28.0% increase from 3 ppm to 5 ppm, 12.0% from 5 ppm to 10 ppm, and 7.0% from 10 ppm to 2%, with the steepest change between 3 ppm and 5 ppm. Final body widths decreased progressively with increasing concentration (panel c), from 3.05–3.12 mm initially to lower widths at higher concentrations, dropping to ∼1.23 mm at 2%, with the sharpest decline between 3 ppm and 5 ppm. Blastema formation was substantial at 3 ppm (∼70%), peaked at 5 ppm, occurred variably at 10 ppm, and was absent at 2% (panel d). At low concentrations (3 ppm), oxidative stress produced considerable but not maximal blastema formation. Only the intermediate concentration (5 ppm) produces an optimal balance, enough injury to trigger regenerative signaling without exceeding the organism’s capacity to recover. Detailed measurements for individual planaria are provided in Supplementary Table S1.

Exposure to H_2_O_2_ produced measurable shifts in planarian zone occupancy over the 28-day conditioning period (Figure 2). Zone A (center) occupancy increased steadily across days, rising from 0.09 ± 0.022 pre-exposure on Day 0 to 0.55 ± 0.012 by Day 28; post-exposure values were slightly lower at each time point. Zone B (annulus) showed intermediate baseline occupancy that was consistently higher post-exposure, increasing from 0.205 ± 0.015 pre-exposure on Day 0 to 0.41 ± 0.011 by Day 28. Consistently, Zone C (outer ring) occupancy decreased over time, with post-exposure proportions starting at 0.70 ± 0.022 on Day 0 and dropping to ∼0.03 by Day 28; post-exposure values followed a steeper decline, reaching 0.015 ± 0.009 by Day 28. Detailed values for all time points and individual zones are provided in Supplementary Table S2. Across all zones, post-exposure values trended lower for Zones A and C but higher for Zone B, with the strongest divergence in the peripheral Zone C. Results confirm that planaria increasingly concentrated in central zones over repeated conditioning sessions, with H_2_O_2_ exposure producing zone-specific shifts, reductions in A and C, but consistent increases in B, most evident in the outer and annular regions.

Paired comparisons confirmed that post-exposure zone occupancy differed significantly from pre-exposure values in most cases (Supplementary Table S3). Zone A showed nonsignificant differences through Day 16 (p ≥ 0.05), but from Day 19 onward pre-versus post-exposure values diverged significantly (p < 0.05), with effect sizes up to d = 6.67 at Day 28. Zone B exhibited significant increases in occupancy at all probe sessions (p < 0.05), with large effect sizes throughout (d = 5.58–10.46). Zone C showed significant decreases across nearly all days (p < 0.05), with stronger effects in the early decline phase (d = 4.42–6.06) and smaller but still significant differences after Day 26 (d = 2.5–2.78).

## Discussion

This study demonstrates that planaria can acquire and maintain conditioned place preferences under a food-reinforced operant conditioning system with light as the conditioned stimulus, and that acute oxidative stress via H_2_O_2_ disrupts these learned behaviors. During the pre-exposure training phase, the progressive increase in Zone A occupancy over the 28-day period indicates successful development and retention of conditioned responses, with peripheral Zone C occupancy declining. This baseline learning confirms that planaria established stable conditioned behaviors prior to H_2_O_2_ treatment, consistent with prior literature on operant conditioning in invertebrates.

Acute H_2_O_2_ exposure produced concentration-dependent effects specifically affecting the expression of conditioned behavior. Moderate concentrations (3–5 ppm) induced prolonged but recoverable erratic movement, reflecting temporary neural and physiological stress without compromising overall viability, whereas higher concentrations (10 ppm and 2%) caused rapid spasmodic movements and severe morphological contraction, reflecting cytotoxicity but exceeding the range relevant for conditioned behavior analysis.

Analysis of post-exposure zone occupancy revealed zone-specific effects. Zone A occupancy remained stable through Day 16, with significant reductions emerging only from Day 19 onward, indicating that the precision of conditioned responses decreases while underlying memory remains intact. Zone B consistently increased in occupancy post-exposure, reflecting that planaria retain the capability for memory, but with less precise localization, as more flatworms occupy Zone B rather than the central Zone A. Zone C occupancy decreased after H_2_O_2_ exposure, consistent with disruption of peripheral, non-reinforced behaviors rather than loss of learned memory. The reduction of Zone A occupancy and increase of Zone B occupancy supports the Interpretation that oxidative stress results in the reduction of the quality of memory post exposure, but that memory capability is still present.

## Conclusion

This study establishes planarians as a tractable model for investigating how oxidative stress influences the relationship between neuronal regeneration and memory. By pairing operant conditioning with controlled hydrogen peroxide exposure, we demonstrated that while planarians retain the capacity to acquire and express conditioned behaviors following regeneration, the precision and retention of these behaviors are measurably diminished. Through determining the optimal range for H_2_O_2_, we further established criteria for later projects dependent on similar use cases to our study. As the first properly documented paper regarding this issue, we aim to use this mini study as an advancement toward academia. The observed reduction of Zone A occupancy and increased occupancy in Zone B highlights that the baseline capacity for memory itself remained constant. However, the quantitative measurement of how strong memory was, or in this case the synaptic connection in the circuit, was not as strong as before. While not a large difference, this discrepancy took place in a model organism, planaria, representing, as mentioned prior, a circuit. When considering the full context of the brain, this finding is significant. These findings reinforce that neurodegenerative disorders such as ALS–FTD cannot be addressed solely through strategies that promote neuronal regrowth. Just as planarians regained structural viability but expressed less precise behavioral memory, patients with neurodegeneration may require interventions that support both structural repair and functional integrity of neural circuits. By bridging behavioral conditioning with regenerative biology, this work highlights the importance of assessing not only whether neurons return, but also how well regenerated circuits preserve and execute learned information.

## Limitations and Future Directions

This study has several limitations that should be considered. First, only a single acute concentration of hydrogen peroxide (5 ppm) was used. Although this dose was selected as reliably sublethal, it does not define how different concentrations of oxidative stress might influence regeneration or learned behavior. Future experiments should test a wider range of doses to determine thresholds for survival, stress, and recovery.

Second, memory was evaluated only through light–place conditioning, which required worms to overcome their innate negative phototaxis. While this provided a clear measure of learned preference, it does not address other forms of behavior such as chemotaxis or vibration response. Adding additional behavioral tests would help determine whether oxidative stress has similar effects across different sensory and motor systems.

Third, oxidative stress was modeled only by acute application of hydrogen peroxide. This approach does not reflect the full range of reactive oxygen species or the effects of long-term exposure, which are common in disease states. Comparing acute and chronic exposures, or using other sources of oxidative stress, would provide a more complete picture of how neurons and behavior respond.

Finally, the recovery period was limited to 14 days. This timeframe was sufficient to observe short-term recovery, but it remains unknown whether the restored behavior is stable over longer periods or whether later deficits may appear. Extending the recovery window and pairing behavior with cellular measures of regeneration would strengthen the conclusions.

Even with these limitations, the results provide a starting point for future studies to test targeted interventions. Compounds with antioxidant or neuroprotective effects, such as curcumin or andrographolide, could be applied during or after injury to accelerate neurogenesis rather than relying solely on natural recovery. Such approaches may alter the quality of memory following regeneration and could reveal whether pharmacological support improves memory retention after oxidative stress.

## Acknowledgments

We would like to express our sincere gratitude to Dr. Xiuxia Du for her role as a corresponding author and mentorship through the final stage of development regarding this paper. We also extend our appreciation to the North Carolina School of Science and Mathematics Department of Sciences for providing access to laboratory facilities, equipment, and faculty supervision that ensured adherence to safety protocols. We further thank the NCSSM Biology and Chemistry departments’ teaching assistants and faculty for their contributions to quality control and ensuring accuracy of experimental protocols.

